# BREADR: An R Package for the Bayesian Estimation of Genetic Relatedness from Low-coverage Genotype Data

**DOI:** 10.1101/2023.04.17.537144

**Authors:** Adam B Rohrlach, Jonathan Tuke, Divyaratan Popli, Wolfgang Haak

## Abstract

Robust and reliable estimates of how individuals are biologically related to each other are a key source of information when reconstructing pedigrees. In combination with contextual data, reconstructed pedigrees can be used to infer possible kinship practices in prehistoric populations. However, standard methods to estimate biological relatedness from genome sequence data cannot be applied to low coverage sequence data, such as are common in ancient DNA (aDNA) studies. Critically, a statistically robust method for assessing and quantifying the confidence of a classification of a specific degree of relatedness for a pair of individuals, using low coverage genome data, is lacking.

In this paper we present the R-package BREADR (Biological RElatedness from Ancient DNA in R), which leverages the so-called pairwise mismatch rate, calculated on optimally-thinned genome-wide pseudo-haploid sequence data, to estimate genetic relatedness up to the second degree, assuming an underlying binomial distribution. BREADR also returns a posterior probability for each degree of relatedness, from identical twins/same individual, first-degree, second-degree or “unrelated” pairs, allowing researchers to quantify and report the uncertainty, even for particularly low-coverage data. We show that this method accurately recovers degrees of relatedness for sequence data with coverage as low as 0.04*X* using simulated data, and then compare the performance of BREADR on empirical data from Bronze Age Iberian human sequence data. The BREADR package is designed for pseudo-haploid genotype data, common in aDNA studies.

## 1 Introduction

An important quality control step in ancient DNA (aDNA) studies is to use estimates of biological relatedness to see if sequence data may come from different skeletal elements or tissues from the same individual, and can hence be merged. Secondly, it is often used to exclude all but one of a group of closely-related individuals for reducing bias in downstream genetic analyses which rely on independent samples of the allele frequency distribution, and thus could lead to biased results. Recently, due to methodological improvements in sampling, DNA extraction, library preparation and capture techniques, archaeogenetics studies have also begun to explore focused regional studies, aimed at investigating kinship and social organisation in groups of burials or even entire graveyards/burial grounds ([10, 18, 1, 24, 17, 20]). Robust estimates of biological relatedness are a key source of information in concert with archaeological and anthropological lines of evidence (among others) for inferring kinship practices in past societies.

Diploid genomes from sequenced individuals are made up of segments of inherited DNA from parent generations, and when inspected to uncover regions that are identical by descent, can be used to infer close genetic relationships. Modern methods for investigating the degrees of genetic relatedness usually require diploid, and even phased, sequence data ([3]). However, due to post-mortem DNA damage and degradation, the coverage of human aDNA sequence data can be extremely low and thus impossible to phase ([12]). Exciting new methods that employ imputation as a pre-processing step ([16]) also require relatively high coverage data (for aDNA), a situation which in, for example, whole-cemetery analyses, is extremely unlikely to be the case for all individuals.

Many methods exist for estimating the degree of genetic relatedness between pairs of individuals for low-coverage aDNA studies. These different methods, based on different types of estimators, have different assumptions and minimum-date requirements, and a comparison of their performance, and their strengths and weaknesses is beyond the scope of this paper. However, for a complete comparison of these software packages, we direct the reader to marsh2023inferring.

Of these methods, we identify READ (Relationship Estimation from Ancient DNA ([11])) as the most similar, peer-reviewed method for estimating genetic relatedness for aDNA, to BREADR. READ performed extremely well when compared to existing genetic relatedness software ([8]), and so will serve as a perfect comparison for the performance of the hard classifications of degrees of relatedness of BREADR, but critically, also for measuring and reporting the uncertainty in these assignments.

Critically, where READ produces a Z-score for whether or not a pair of individuals can be “more or less” related (*Z*_lower_ and *Z*_upper_ respectively), this is calculated by subtracting the midpoint between the expected PMR for the lower and upper classes of relatedness (divided by the jackknife-estimated standard deviation). This approach treats the density of the PMR for each related class as uniform, and non-overlapping, with the expected PMR lying precisely at the midpoint between these boundaries. BREADR instead assumes a binomial distribution for the distribution of the PMR values, and using the associated probability density functions, returns not just a hard classification (the most “likely” genetic relationship between each pair of individuals) but also the posterior probabilities of all possible genetic relationships in detailed diagnostic plots. With this complete information, researchers are able to better make decisions on pairwise relatedness, informed by rigorous statistical calculations, with measures of uncertainty when considering alternative genetic relationships.

We identified that no PMR-based method includes a statistically rigorous measure of classification uncertainty, and that no method includes diagnostic plots for interpretation. Here we present BREADR: a simple-to-use package for the R-statistical software. We compare the performance of BREADR to READ, a popular, field-standard, peer-reviewed method for estimating biological relatedness via the PMR for up to the second-degree from low-coverage aDNA data.

## 2 Comparisons to Existing Software

Most software packages for estimating genetic relatedness from aDNA sequence data leverage some measure of pairwise genetic dissimilarity, often calculated on pseudo-haploid data, where a random call is sampled for each site, to avoid the issue of phasing.

lcMLkin uses genotype likelihoods to estimate identity-by-descent coefficients (*k*_0_, *k*_1_ and *k*_2_), from which the co-ancestry coefficient can be calculated, such that *θ* = *k*_1_*/*4+*k*_2_*/*2 ([6]). This method requires a reference panel to estimate the genotype likelihoods, and for sites to be unlinked. This method is exploratory in nature, and does not return hard classifications of degrees of relatedness, and is (currently) not peer-reviewed.

TKGWV2 instead estimates relatedness via the method of moments estimator from Queller and Goodnight, denoted 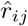. This asymmetric estimator assumes no inbreeding, in the population and that all sites are unlinked ([15]). Hence, TKGWV2 cannot be applied to capture data, or data from populations where inbreeding has occurred. This method returns hard classifications of degree of relatedness based on which expected value of 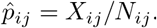 the observed value is closest to.

KIN ([14]) and ancIBD ([16]) instead use the spatial distribution of identical-by-descent segments (IBDs). Like lcMLkin, KIN estimates the identity-by-descent coefficients, using IBDs instead of weighted genotype likelihoods, and can identify degrees of relatedness up to the third degree. Conversely, ancIBD uses imputation to increase the resolution of the method up to the seventh-degree at the cost of requiring relatively high-coverage sequence data.

READ uses the so-called pairwise-mismatch rate (PMR) to estimate the proportion of overlapping sites for which two individuals have non-matching genotype calls ([11]). The PMR was first introduced to overcome the limitations of low-coverage aDNA, however, the publication from Kennett *et al*. did not include a hard-classification method. Where Kennett *et al*. used all of the available overlapping site to estimate the PMR, READ uses a 1MB windowed approach to sample from the distribution of the PMR, but also to overcome the effects of linkage disequilibrium (LD), allowing for the automated hard classification of degrees of relatedness. BREADR, like READ, estimates the PMR, but instead thins the data to overcome the effects of LD. The PMR on this thinned data can now be reasonably expected to follow a binomial distribution, and theoretical expectation of the PMR are derived and compared to the observed value. From this, statistically rigorous measures of uncertainty are derived, and informative diagnostic plots of the degree of related for pairs of individuals can be produced. Finally, although BREADR only classifies degrees of related up to the second-degree, we also allow for tests of any degree less than the second-degree to be formally tested when considering potential pedigree reconstructions.

### 2.1 Overview of the Paper

The paper is structured as follows. First we begin by briefly describing the theory that allowed for the development of the PMR (Section 3), followed by a description of our implementation of the BREADR method. We then describe the two data sets: one simulated and one empirical (Section 4). Using the simulated data, we provide a technical description of the functionality of the implemented functions, focusing on their interpretation and comparing the results to those obtained from READ (Section 5). Following this we then analyse empirical data to assess the performance of BREADR, and to again compare our results to those obtained from READ (Section 5.6). Finally we discuss the strengths and limitations of the methods implemented in the BREADR R package (Section 6).

## 3 Methodological Background

The quantification of degrees of relatedness have existed since the early days of population genetics in the early twentieth century via the coefficient of relationship, denoted *r*_*ij*_ for individuals *i* and *j*, ([23]). To calculate *r*_*ij*_, a pedigree was required, allowing a path coefficient between *A* (an ancestor), and *O* (an offspring), separated by *n* generations, to be defined as

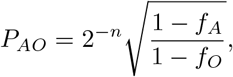

where *f*_*A*_ and *f*_*O*_ are inbreeding coefficients.

The coefficient of relationship *r*_*ij*_ could then be calculated, via a most recent common ancestor of both

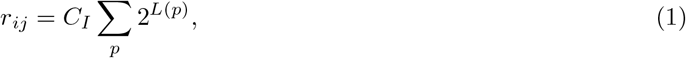

where *L*(*p*) is the length of path *p*, and we assume that the inbreeding coefficient is constant for the entire pedigree. Note that this summation occurs over all possible paths *p* between individuals *i* and *j*, which, for example, for siblings contains two paths via the maternal and paternal line, but for parent and child, only one path exists.

However, as pedigrees are rarely available, and are in fact the very thing researchers often wish to reconstruct, methods for estimating measures of relatedness from genetic markers have been proposed in the last several decades ([7, 15, 5, 19]). Of special note though was the kinship coefficient, denoted *ϕ*_*ij*_ ([2]), which, under some assumptions such as limited inbreeding, is related to Wright’s coefficient of relationship via

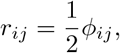

providing a direct relationship between the degrees of relatedness from the pedigree, and the expected kinship coefficient *ϕ*_*ij*_ (see Table 1).

**Table 1:** Values of the coefficient of relatedness and the kinship coefficient for different pedigree relationships up to the second-degree, assuming that *C*_*I*_ = 1.

| Relationship | Degree | Coefficient of Relationship | Kinship Coefficient |
| --- | --- | --- | --- |
| Same Individual/Twins | 0 | $2^0 = 1$ | $1/2$ |
| Parent-Child | 1 | $2^{-1} = 1/2$ | $1/4$ |
| Sibling-Sibling | 1 | $2 \cdot 2^{-2} = 1/2$ | $1/4$ |
| Grandparent-Grandchild | 2 | $2^{-2} = 1/4$ | $1/8$ |
| Aunt/Uncle-Niece/Nephew | 2 | $2 \cdot 2^{-3} = 1/4$ | $1/8$ |
| Half-Siblings | 2 | $2^{-2} = 1/4$ | $1/8$ |

An interpretation of 1*−ϕ*_*ij*_ is that it is the probability that a randomly sampled allele from each individual is not identical by descent. Assuming that alleles are not under linkage disequilibrium, it follows that we may estimate this value from genome-wide data for a pair of individuals. However, limitations due to low numbers of sampled individuals and low genome coverage have been found to complicate and bias estimating these values, making genetic relatedness in aDNA studies difficult ([21, 22]).

The pairwise mismatch rate (PMR) was first introduced in the aDNA literature by Kennett *et al*., and later refined and presented as a software package by Monroy Kuhn *et al*. ([4, *11]) to overcome these issues. The PMR can be used to estimate the kinship coefficient, which assuming that we can account for the inbreeding coefficient C*_*I*_, can be used to estimate the degree of relatedness, up to some reasonable resolution, and hence we may gain insights into the pedigree joining many individuals (to a certain resolution).

### 3.1 The Statistical Model

Consider two individuals, *i* and *j*, with *N*_*ij*_ overlapping sites without missingness, where sites are thinned (default value 1×10^5^) such that the effects of linkage disequilibrium are reduced. We then look at the number of pseudo-haploid genotype calls that *do not* match, denoted *X*_*ij*_. Hence we have that *X*_*ij*_ *∼ Bin*(*N*_*ij*_, *p*_*ij*_), and that the maximum likelihood estimator for *p*_*ij*_ is 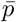.

Once all PMRs are calculated, we must account for background relatedness, which can be thought of as the expected PMR for a pair of unrelated individuals. Many choices exist for this value, but assuming that a sample is made up of mostly unrelated pairs (to the second degree), then the median PMR, denoted 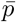 will be a reliable estimate ([11]). However, we also allow 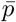 to be a user-supplied parameter, which may be known from previous studies, or can be varied for sensitivity analyses. Note that 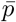 can be thought of as an estimate of *C*_*I*_ from Equation 1.

Borrowing from the insights of READ, we now define the expected mean PMR for a relationship of degree *k* = 0, 1, 2 to be

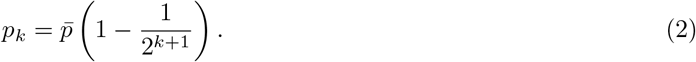

Hence, if we assume that the degree of relatedness for individuals *i* and *j* is truly of the *k*^th^ degree, then

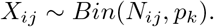

Hence, the likelihood function for relatedness degree *k*, for individuals *i* and *j*, is

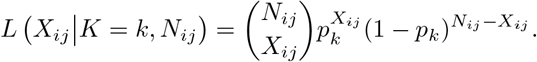

If we let *k* = *∞* be the case that two individuals are “unrelated” (*i*.*e*. more than second-degree related), but then we also have that

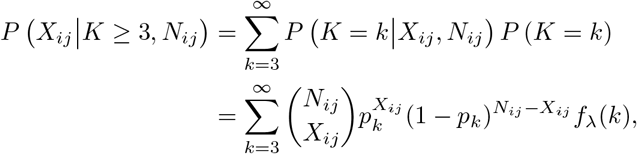

where *f*_*λ*_(*k*) is the 3-truncated Poisson distribution of the form

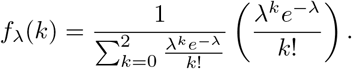

We choose the 3-truncated Poisson distribution with *λ* = 10 as it represents well the unlikelihood of individuals always being closely related once they are more than second-degree related, but also captures the diminishing probabilities of being too distantly-related due to the finite size of populations.

It is then possible to calculate the normalised posterior probabilities of individuals *i* and *j* being *k*-degree related as

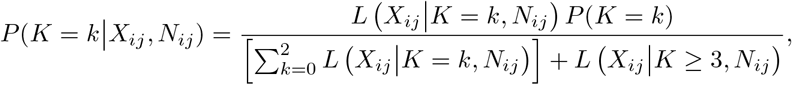

for which the denominator, by construction, equals one. Hence,

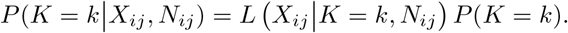

It is not clear what the prior probabilities for the degrees of relatedness, from same/twin to second-degree related, should be, and so these are given as uniform by default. However, since these are user inputs, these can be explored via sensitivity analyses to test if the prior probabilities are driving the relatedness classifications.

Finally, while we do not include third-degree relationships in the possible classifications, we do include a method for statistical test for retaining or rejecting the possibility of a *k*^th^-degree relationship, for 0 ≤ *k* ≤ 10. We simply perform a binomial test for the observed number of mismatches *X*_*ij*_, and the expected probability of mismatches, as defined in Equation 2, returning a two-sided p-value and the estimated (non-integer) degree of relatedness, found by setting *p*_*k*_ = *p*_*ij*_ and 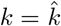into Equation 2, and solving for *k, i*.*e*.,

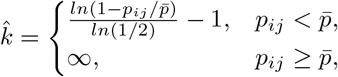

where *∞* indicates “unrelated”.

## 4 Data Sets

To test the performance of BREADR, we performed analyses on simulated data as described in [14]^1^, and on published empirical data from a population genetic and relatedness study by villalba2022kinship^2^, and compared our results to those obtained from READ.

### 4.1 Description of Simulated Data

We simulated genotype data from a pedigree of individuals as shown in Figure 1. This pedigree has 2 genetically identical pairs of individuals, 19 first-degree relatives, 12 second-degree relatives (2 of which are half-siblings) and 87 unrelated pairs of individuals. The pedigree also contains a further 9, 6 and 1 third-, fourth- and fifth-degree relatives, respectively, although our method does not attempt to directly identify these deeper relationships.

**Figure 1:**
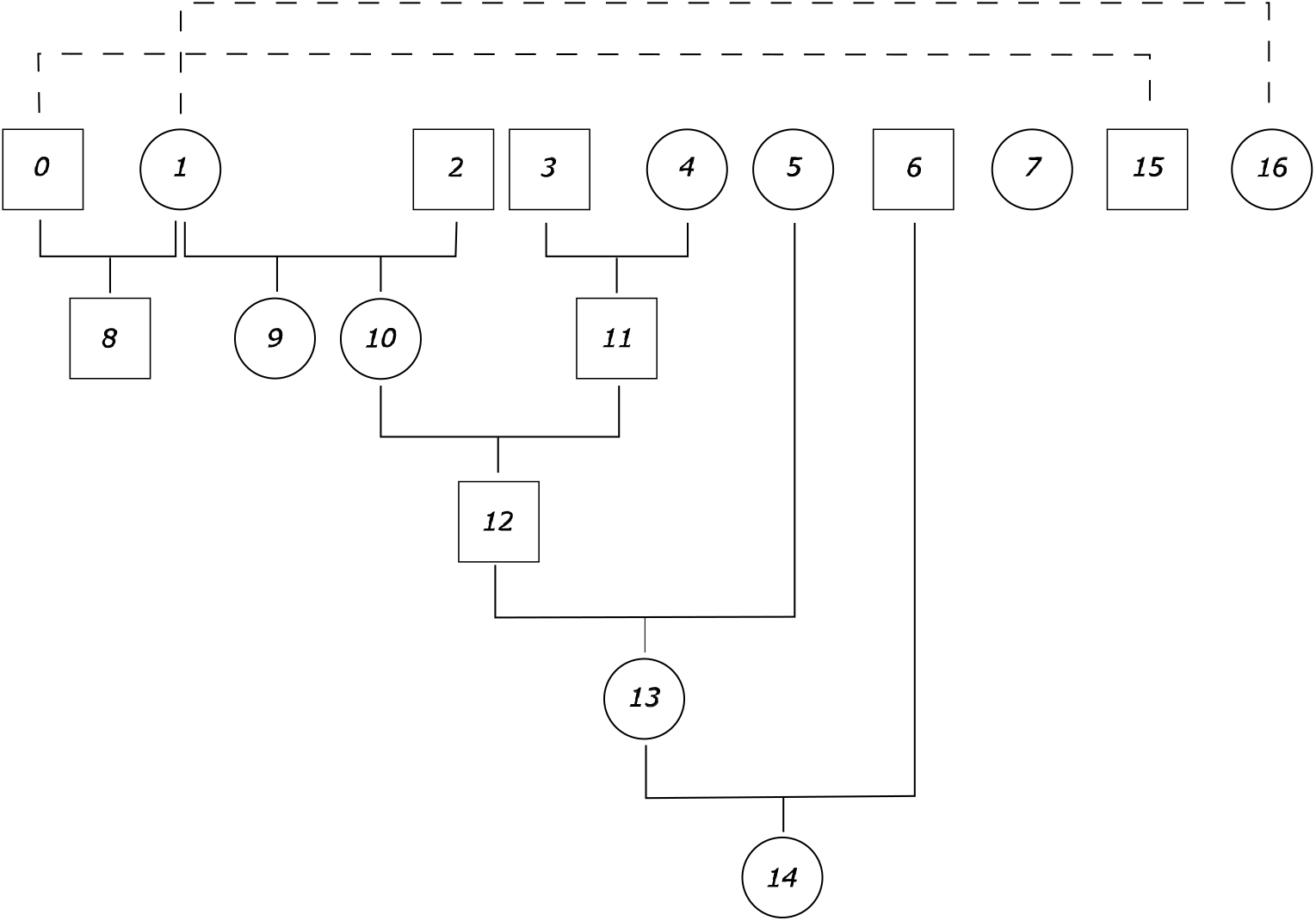
Pedigree for the simulated data. Individuals 0 and 8, and 1 and 9, are genetically identical, respectively. Females are represented by circles, males are represented by squares.

The simulated data resulted in genotypes being called for 29,767,814 segregating sites, at an average of 0.04X coverage.

### 4.2 Description of the La Almoloya data set

To test the performance of BREADR on empirical data, we analyse a pseudo-haploid Eigenstrat data set ([13]) generated from prehistoric individuals who were buried at the Early Bronze Age site La Almoloya in today’s Spain ([20]). This data was generated from 1240k capture ([9]) data using pileupcaller^3^ with mapping and sequence quality filtering parameters -q 30 and -Q30, and the –randomHaploid flag. The data consists of 68 individuals, and included 13 pairs of first-degree individuals, 10 pairs of second-degree related individuals, and 2255 unrelated pairs of individuals. The authors identified these relationships using a suite of information including: genetic relatedness estimated by READ, lcMLkin, shared runs of homozygosity, uniparentally-inherited markers from the mitochondrial genome and the Y chromosome, as well as archaeological and anthropological contextual information, such as age at death and stratigraphy. For details on the bionformatic processing of the raw sequence data, and for data quality statistics such as coverage, we refer the reader to the original study ([20]).

## 5 Package Structure and Usage applied to Simulated Data

The package BREADR is available under the MIT license from the Comprehensive R Archive Network (CRAN) at https://CRAN.R-project.org/package=BREADR. The package can also be installed and loaded using the following commands: The package contains the functions for a full analysis of the genetic relatedness for a large number of individuals, and the visualisation of the overall genetic relatedness, as well as pairwise diagnostic plots to make informed pedigree reconstructions.

The raw input data is genotype data called in the Eigenstrat format, and this allows for user pre-processing for data characteristics such as: removing ancient DNA damage, minimum allele frequency cut-offs, random versus majority genotype calls, read and mapping quality scores, etc.

An analysis begins with a processing step where genotype data is converted into count data for each pair of individuals. As this is the most computationally intensive step in the analysis, all downstream analyses are performed on the output of this first step. From this data we calculate the genetic relatedness statistics, calculate posterior probabilities, and make highest posterior probability assignments, for the degrees of genetic relatedness.

Lastly, we provide functions for visualising the genetic relatedness estimates and classifications, and for performing adhoc tests for user defined degrees of relatedness via a binomial test.

We ran the algorithm with the default thinning parameter of 1 × 10^5^, minimum number of overlapping sites of 500, a uniform prior for the class classifications, and the median used to estimate the background relatedness. Overall, we correctly assigned 129/136 (94.85%) of the pairwise genetic relationships, indicating a relatively high degree of accuracy.

BREADR correctly classified 2/2 of the genetically identical pairs with high posterior probability (one to machine precision), and 19/19 of the first-degree related pairs (minimum posterior probability 0.993). BREADR correctly identified 10/12 of the second degree pairs, and in the two cases where we misclassified the relationships (both times as unrelated) READ also misclassified the second-degree pairs as unrelated. Furthermore, READ misclassified an additional two first-degree related pairs as unrelated.

Finally, BREADR correctly classified 98/103 of the unrelated and third-degree or higher related individuals as “unrelated”. Interestingly, the individuals in the remaining five cases were actually related in the third-degree in the pedigree. This was reflected in the fact that, even though there was strong evidence for classifying these individuals as second-degree related compared to unrelated, the binomial tests for third-degree were all non-significant. We note that READ did not misclassify these five relationships, and returned an absolute Z-score less than 2 in 4/5 cases, indicating that a closer relationship than unrelated was also possible.

### 5.1 Preprocessing the Eigenstrat data for analysis

All downstream functions in the BREADR package require preprocessing of an Eigenstrat “trio”: so-called ind, snp and geno files. During this first step, we take the list of sites on the genome of interest, with the chromosome name and (integer) site position, and the pseudo-haploid genotype calls for all individuals (of interest). Then, for each pair, we take only sites which have overlapping, non-missing calls, and are at least some user-defined number of positions apart (within chromosomes). Using these site positions, we record the number of filtered, overlapping sites, as well as the number of mismatches per pair.

To generate these required genetic pairwise comparisons for all individuals, users apply the `processEigenstrat` function. This function requires three input string parameters: the paths to the *ind, geno* and *snp* files that comprise an Eigenstrat trio. At this stage of pre-processing, users have four additional parameters that can be set, and cannot be reset downstream.

First, the *filter length* parameter (default 10^5^) allows the user to define the minimum number of positions between sites that can be considered. By removing sites which are close to one another, the assumption of independence for each site-wise comparison is best attained by lowering or removing the effects of linkage disequilibrium. Second, the *pop pattern* parameter (default exclude nothing) allows the user to define a set of population names so that only a subset of the individuals is compared. Third, the *filter deam* parameter (default NO) filters C-¿T or G-¿A SNPs from the possible list of sites if the potential effects of post-mortem deamination have not been accounted for in the genotyping process. Last, since pre-processing is the most computationally expensive step of the analysis, the *outfile* parameter (default NO) is a path-string that allows the user to automatically save the post-processed data as a TSV file.

The resulting (see Figure 2) tibble has four columns: the names of the samples/individuals which were compared (*pair*), the number of overlapping SNPs (*nsnps*) per pair, the number of overlapping sites for which the pair did not match (*mismatch*) and the pairwise-mismatch rate (*pmr*).

**Figure 2:**
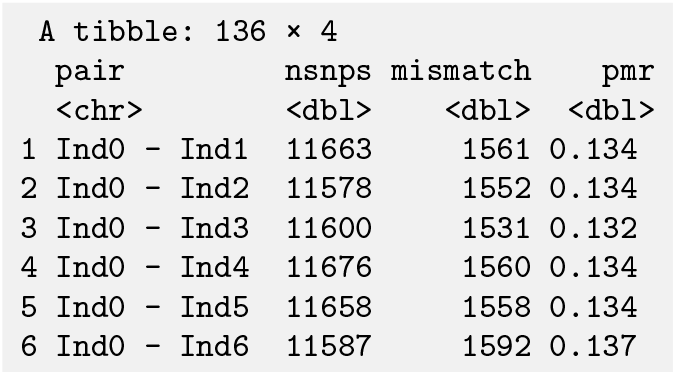
The first 6 rows of the tibble produced by running processEigenstrat() on the simulated data set.

### 5.2 Analysing the post-processed data

Following the pre-processing step, the `callRelatedness` function calls a relationship of Same_Twins, First_Degree, Second_Degree or Unrelated, with additional information, from the output of the `processEigenstrat` function. Additional parameters can be set to allow the user to customise the analysis.

The *class_prior* parameter (default Uniform) defines the prior probabilities of each relatedness class. The *median* parameter defines what the background relatedness should be (default is the median from the filtered estimates). This can be either: (a) estimated directly from the median PMR from the data, (b) a single value (which is useful for sensitivity analyses when the user is uncertain), or (c) a vector of values equal to the number of rows in the input tibble (which can be useful if different populations have different background relatedness levels). The *median co* parameter (default 500) defines the minimum number of overlapping SNPs a pair of individuals must share for their PMR to be used in estimating the median PMR, if the user has elected to use the median PMR. Finally, the *filter n* parameter (default 1) is used to simply remove any pairs of individuals from the analysis if they share less overlapping SNPs than this value.

The resulting tibble (see Figure 3) has an additional 9 columns: the row number (*pair* which is useful for additional functions, the highest-posterior genetic relationship (*relationship*), the standard error of the estimate of the PMR (*sd*), the median used in the calculations (*med*), and the normalised posterior probabilities for each of the four relatedness categories (*Same_Twins, First_Degree, Second_Degree* and *Unrelated*)).

**Figure 3:**
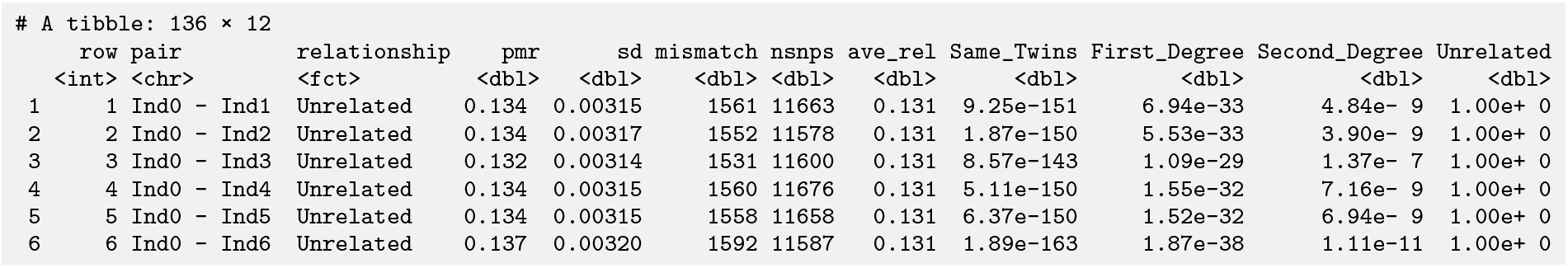
The first 6 rows of the tibble produced by running `callRelatedness()` on the simulated data set as in Figure 2

### 5.3 Visually interpreting results

Once posterior probabilities of degrees of relatedness have been calculated, the results can visualised in two ways.

First, an overall picture of the relatedness can be obtained using the plotLOAF() function which plots the first *N* (by default *N* = 50) pairs of individuals, sorted by ascending PMR value (see Figure 4). The shape and colour of the point estimates of the PMR indicate the highest posterior degree of relatedness assigned for each pair, with an associated 95% confidence interval represented by the vertical error bars. The dashed coloured lines indicated the expected PMR for each degree of relatedness (given the background relatedness). Note that if we have *n* individuals, then we will have 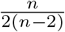 pairs of individuals, which grows factorially as *n* gets larger. Hence plotting all pairs of individuals will be infeasible in many cases, and since the majority of pairs will likely be unrelated, users may only wish to plot the closely related pairs of individuals for brevity. From Figure 4 we can immediately observe that we have two genetically identical individuals, and we also get a scale of the number of first-degree related, second-degree related individuals.

**Figure 4:**
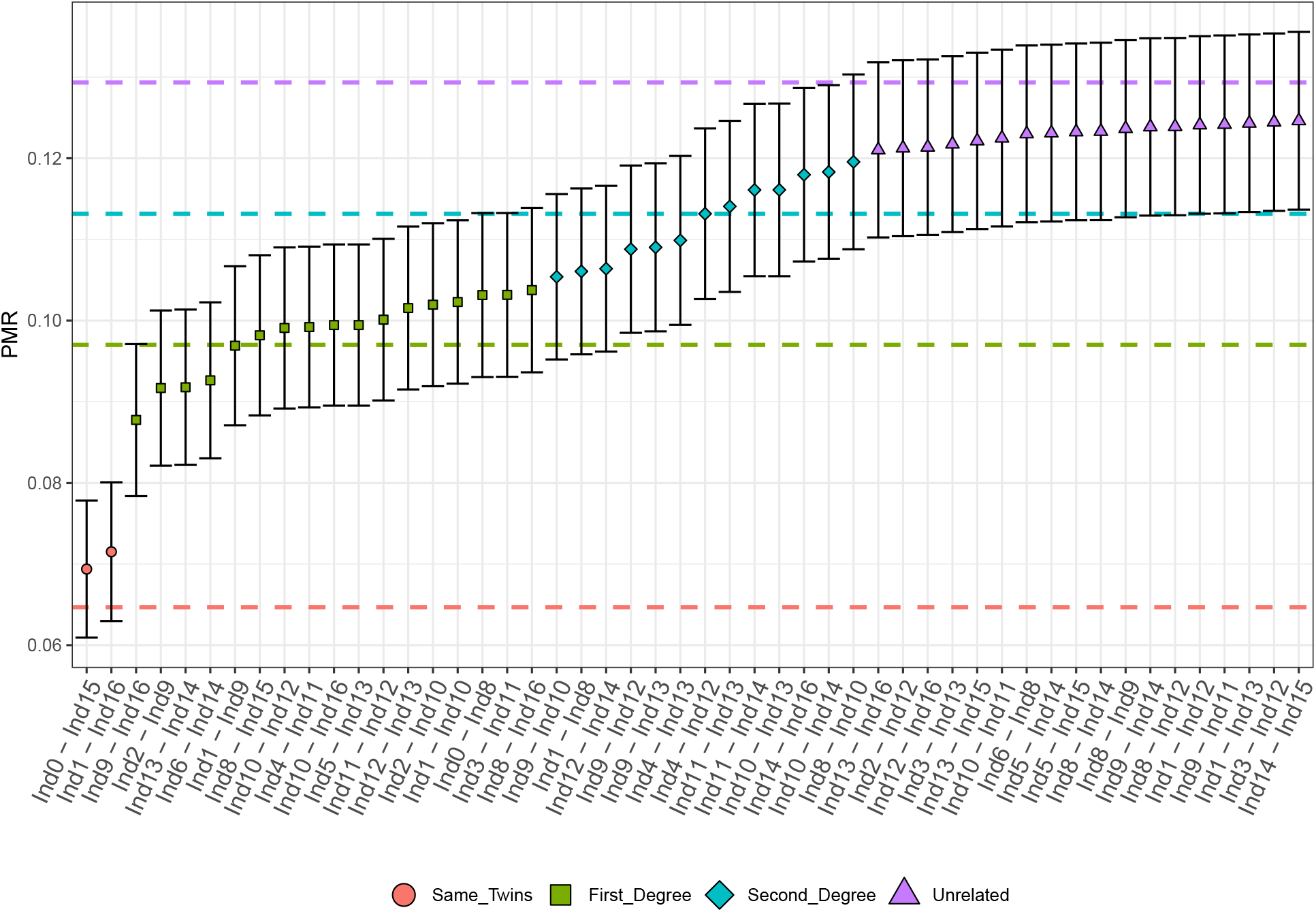
The first 50 ordered pairwise-mismatch rates (PMRs) from the simulated data. Colours indicate classification, dashed lines indicate the expected PMR for each degree of relatedness, and the error bars indicate 2 standard errors.

Second, a diagnostic plot for the analysis of a single pair of individuals can be obtained using the callSLICE() function (see Figure 5. By default, this function produces a two panel plot. The left panel displays the distribution of the PMR for each of the degrees of relatedness (given the number of overlapping sites), as well as the observed PMR (and 95% confidence interval) plotted below these densities. The right panel displays the normalised posterior probabilities for each possible degree of relatedness, both visually and numerically. The user can choose to return just one of these plots, or both. In Figure 5 we misclassified the true second-degree relationship as unrelated, however, the posterior probabilities should lead a researcher to consider a second-degree relationship plausible.

**Figure 5:**
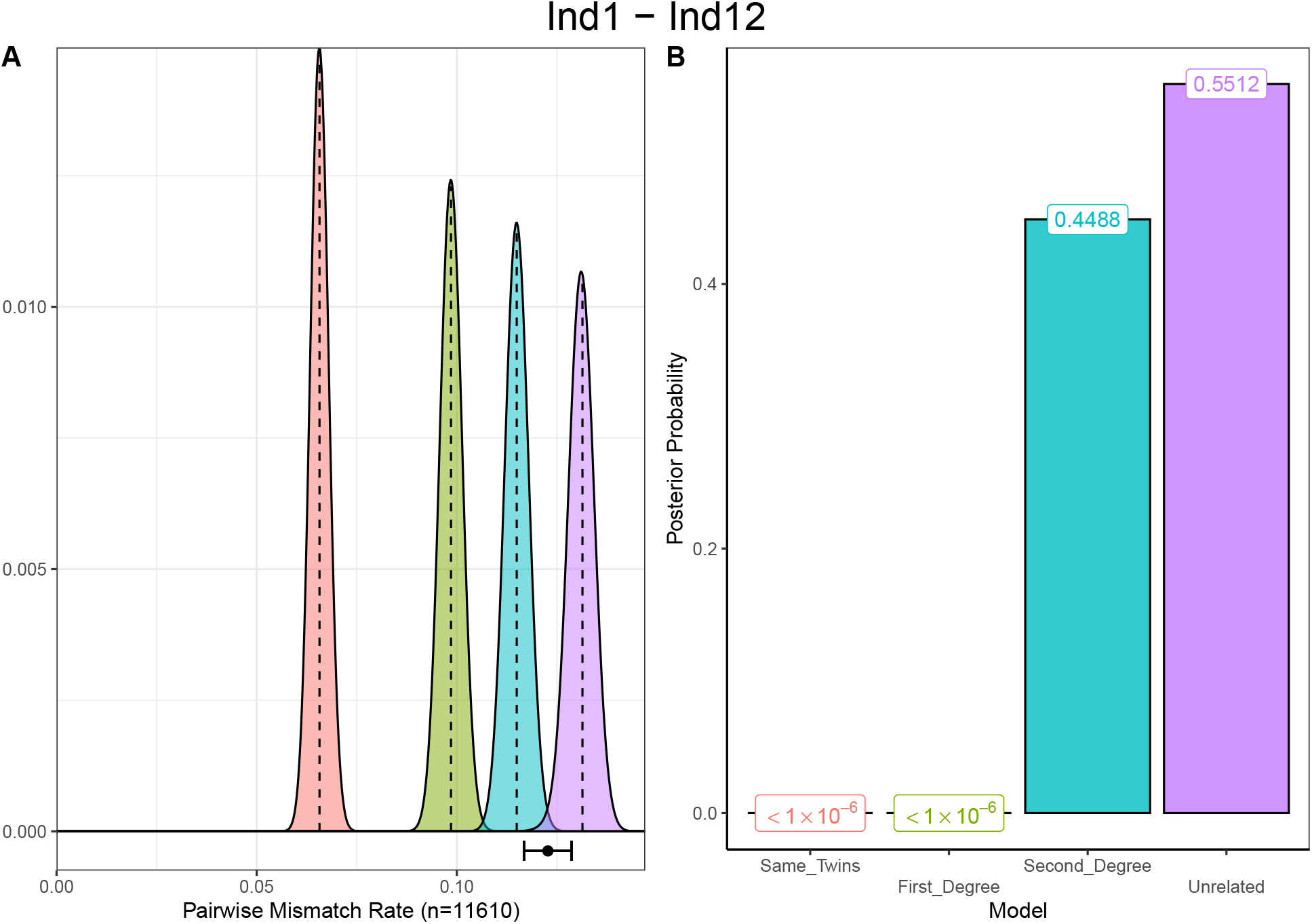
The diagnostic plot for the analysis of the genetic relationship Ind1 and Ind12. Panel A indicates the observed PMR falls between the expected distributions for a second-degree relationship (blue) and an unrelated pair (purple). This is uncertainty is quantified in panel B where the normalised posterior probability assigned to a second-degree relationship is 0.4488, and 0.5512 for unrelated.

Finally, the savesSLICES() function allows the user to save all possible pairwise diagnostic plots (as produced by plotSLICE()) as PDFs to an output folder.

### 5.4 Testing for departure from refined degrees of relatedness

The resolution of the BREADR method only allows degrees of relatedness of up to the second degree to be assigned. However, once can in principle test to see if the observed PMR is consistent with *any* degree of relatedness using a simple binomial test.

The test_degree() function allows the user to test any degree of relatedness up to the tenth degree. If the verbose option is set to TRUE, then all of the information about the binomial test is displayed (see Figure 6). From this the null hypothesis, the expected and observed PMR, the estimate of the degree of relatedness, the associated p-value and the decision (at significance level *α* = 0.05) are given. Figure 6 shows a case where we are able to correctly identify that a third-degree relationship is reasonable for Ind13 and Ind16, which was not immediately obvious in just the diagnostic plot (Figure 7).

**Figure 6:**
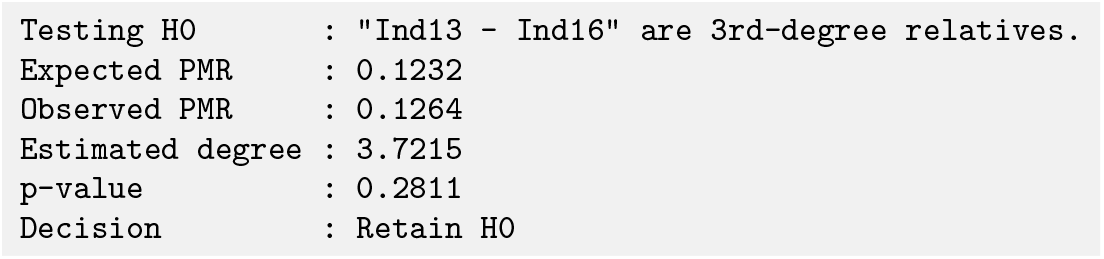
The output from the binomial test for whether the PMR observed for Ind13 and Ind16 is consistent with a third-degree relationship. We correctly classified the relationship as unrelated, as we only report degrees of relatedness to the resolution of the second-degree. However, the binomial test reveals that the observed data is not inconsistent with a third-degree relationship (*p* = 0.2811).

**Figure 7:**
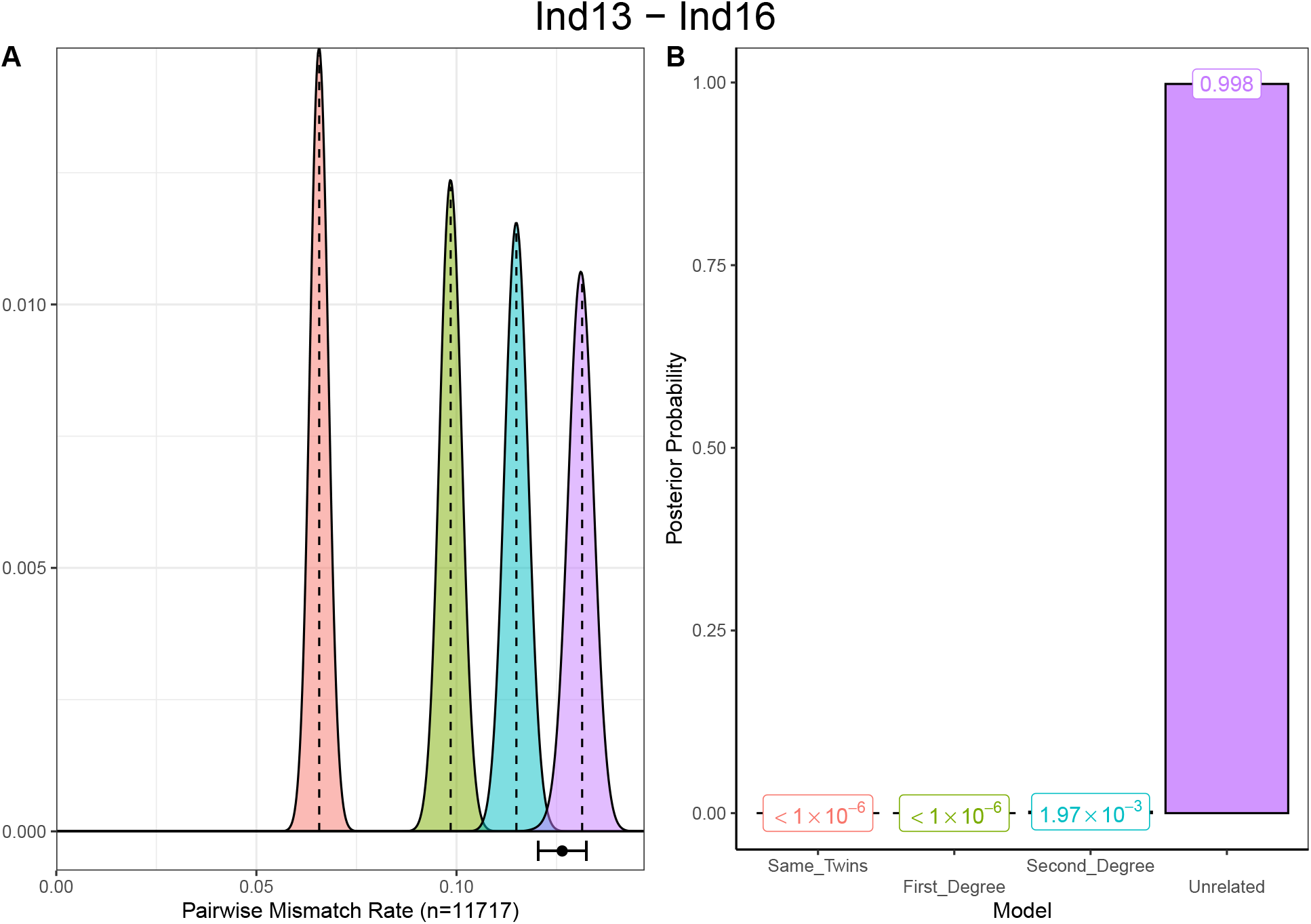
The diagnostic plot for the analysis of the genetic relationship Ind13 and Ind16. Panel A indicates the observed PMR falls between the expected distributions for a second-degree relationship (blue) and an unrelated pair (purple). Unlike for Ind1 and Ind12, this uncertainty is not quantified in panel B, however, the binomial test still reveals the uncertainty in this call.

### 5.5 Comparison of Measures of Uncertainty for BREADR and READ

READ classifies degrees of relatedness by considering the midpoint between the expected normalised PMR values. READ estimates the PMR by partitioning the genome into 1MB windows, and calculating a sample of *n* PMR values, from which a mean (*p*_0_) and standard deviation (*σ*_*p*0_) can be estimated. If the observed *p*_0_ falls between the expected PMR value for degrees of relatedness *k* and *k* + 1, defined in READ as

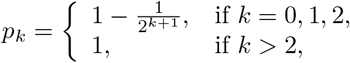

where the midpoint between *p*_*k*_ and *p*_*k*+1_ is

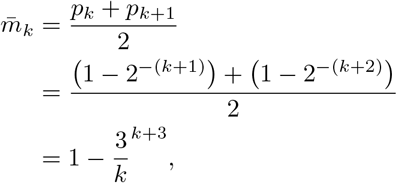

READ then classifies the degrees of relatedness as

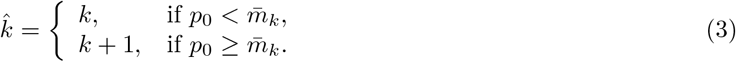

To assess the confidence of the assignment of 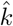 degrees relatedness, compared to degrees 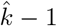 and 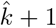, READ then calculates Z-scores of the form

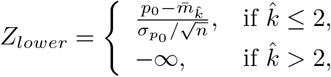

And

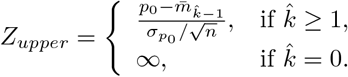

This approach of hard-classifying the degrees of relatedness makes sense, as the midpoint decision criteria in Equation 3 means that the point estimate must be closer to the expected PMR for 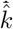. However, using 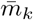 as a cutoff for a Z-score appears arbitrary, especially when one considers sample quality, represented by *N* (the number of overlapping SNPs) in Figure 8, which we use as a proxy for *n*, the number of 1MB windows that yielded PMR values for READ.

**Figure 8:**
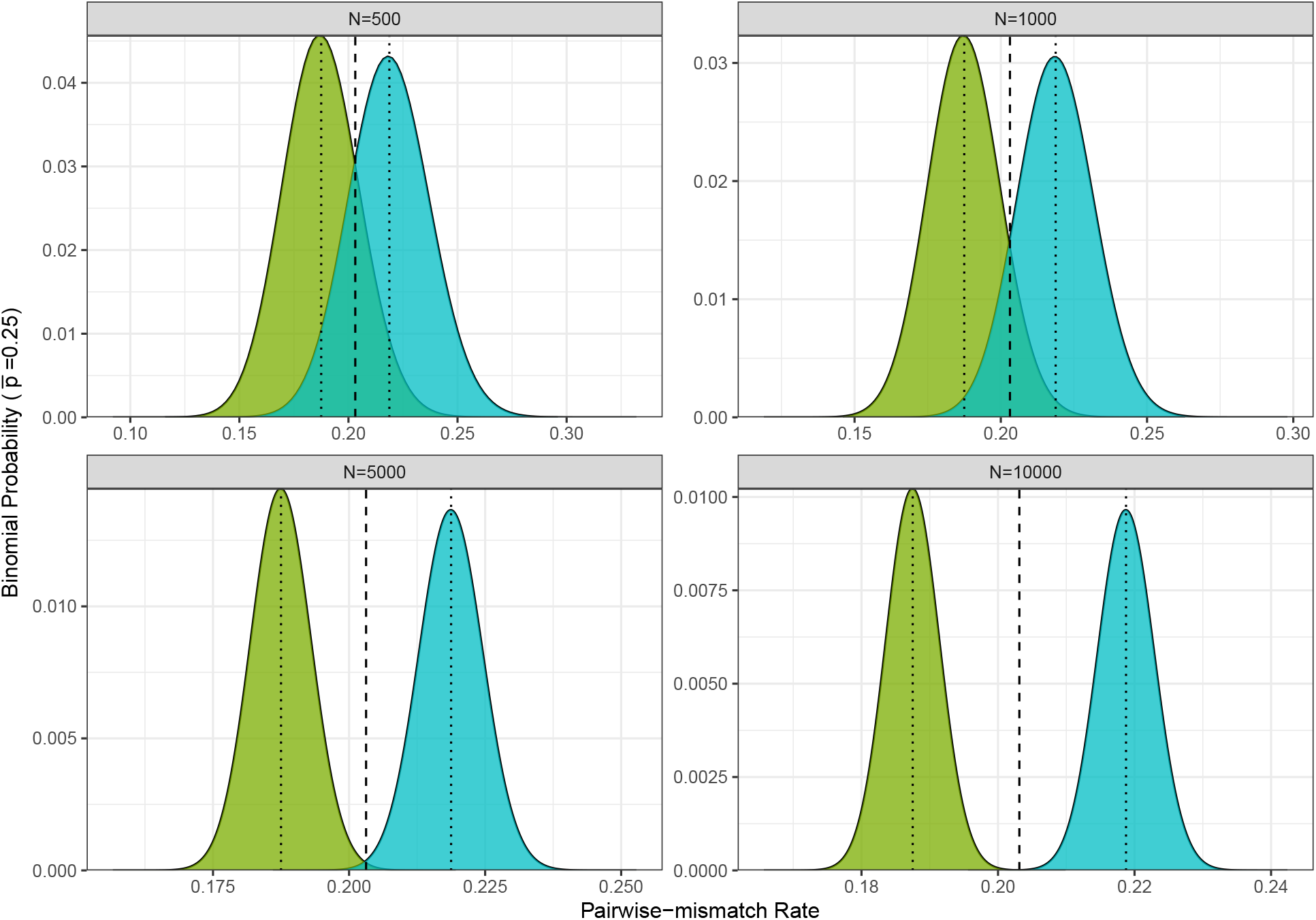
Distributions of expected pairwise mismatch rates (PMR) for degrees of relatedness *k* = 1 (blue) and *k* = 2, with an expected PMR for unrelated individuals of 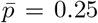. The dotted lines represent the expected PMR for first- and second-degree related individuals, and the dashed line represents the midpoint 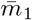 between the expected values.

Clearly, as the number of overlapping SNPs increase, *i*.*e*. sample quality increases, the two distributions separate more. However, in cases where there are low numbers of overlapping SNPs (*N* = 500 and *N* = 1000), a reasonable proportion of the probability density for both degrees of relatedness can be found extended past the midpoint 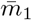. Hence we would expect to find many observed PMR values from first-degree relationships that are greater than 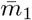. In cases where sample quality is higher, that is *N* is larger (*N* = 5000 and *N* = 10000), 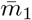 becomes an arbitrarily distant point from both expected PMR values, with very little probability mass associated with the area around 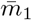, making this point largely probabilistically irrelevant to either distribution.

BREADR quantifies uncertainty differently, and in three ways: the error bars around the point estimate (see Figure 4), the normalised posterior probabilities, and (usually employed in cases where a degree of relatedness greater than 2 might be considered) the binomial test for degrees of relatedness. Our approach has three significant benefits. First, users can consider degrees of relatedness other than one above and below the degrees of relatedness that were assigned for a pair of individuals. Second, instead of the binary decision one can make using a Z-score (for which it is not clear which cut off should be used), users are given a quantitative measure of confidence via the posterior probabilities. Third, our method is statistically rigorous, and does not rely on the distance to an arbitrary, albeit seemingly intuitive, midpoint.

To see the effect of this, we considered cases for the simulated data where READ had identified the correct degree of relatedness, but had also produced |*Z*| < 2, *i*.*e*., that the classification of *k* is uncertain. We found that in the 6/19 (31.57%) of the first-degree relationships, that READ produced |*Z*| < 2, and BREADR returned confident classifications with a mean normalised posterior probability of 0.998. We also found that in 6/12 (50%) that READ produced |*Z*| < 2, where BREADR again returned confident classifications with a mean normalised posterior probability of 0.999. Finally, READ returned that four of the unrelated calls were uncertain, indicating that a second-degree relationship was not unreasonable. In all four of these cases, BREADR also returned low posterior probabilities for “Unrelated” (< 0.135), with higher normalised posterior probabilities for second-degree. Interestingly, in every case, these pairs of individuals were third-degree related, and using the binomial test, we could identify that the observed PMR was not inconsistent with a third-degree relationship (*p* > 0.1874).

Hence, while the hard classifications of degrees of relatedness for READ and BREADR are equally accurate, it can be seen that BREADR describes and quantifies uncertainty in a more statistically rigorous framework, yielding certainty for correct relationship assignments when READ did not.

### 5.6 Empirical data from La Almoloya

We tested our method on the empirical data from the Early Bronze Age site of La Almoloya, in Spain ([20]) using the same parameters as for the simulated data. A plot of the overall relatedness can be found in Figure 9.

**Figure 9:**
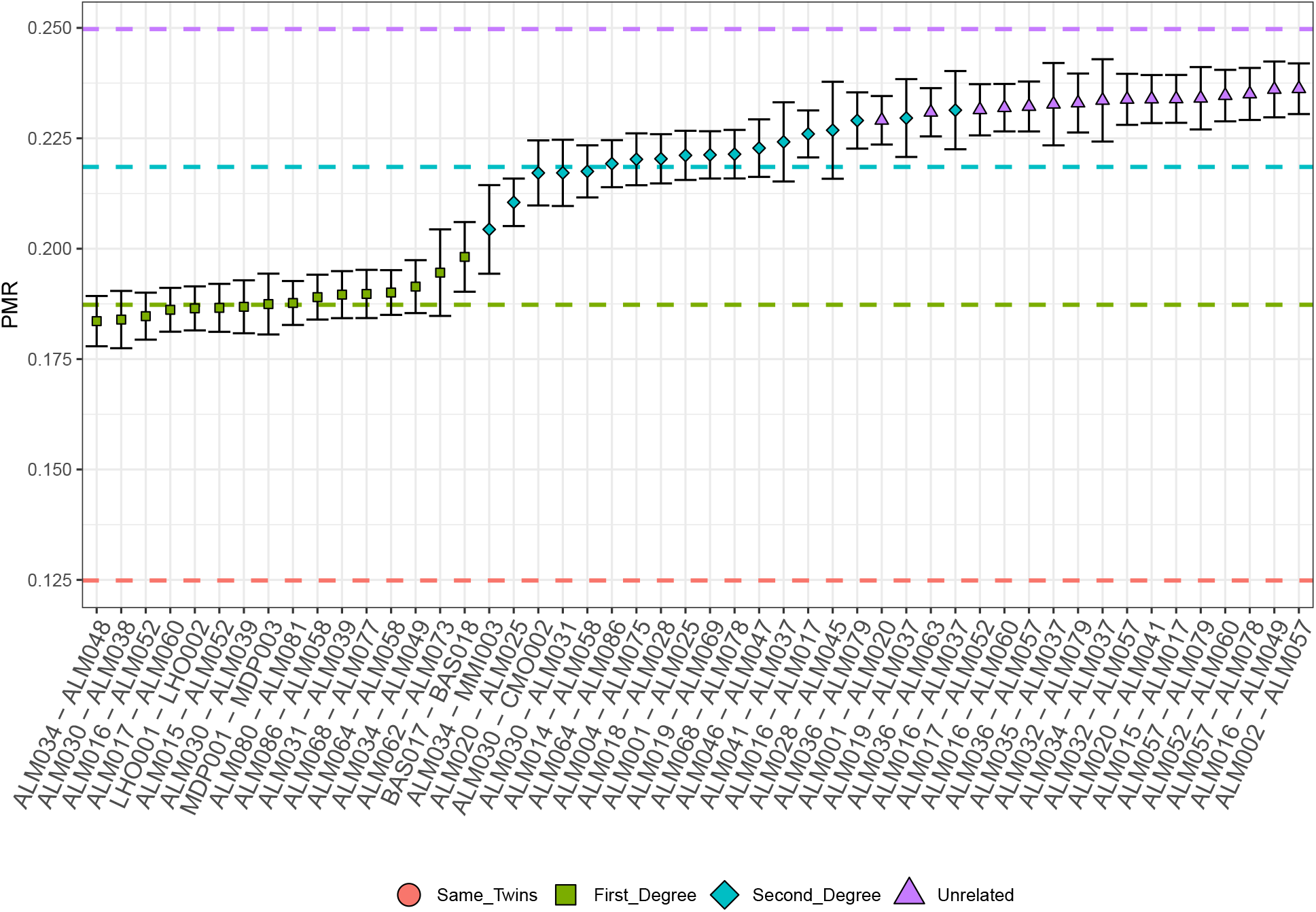
The first 50 ordered pairwise-mismatch rates (PMRs) from the La Almoloya data. Colours indicate classification, dashed lines indicate the expected PMR for each degree of relatedness, and the error bars indicate 2 standard errors.

When we compared our results to READ, we found remarkable consistency with the results from READ. We found that we only disagreed on 2/2278 pairs of individuals: ALM016/ALM017 and ALM057/ALM079.

In both cases, READ classified these relationships as unrelated, whereas our method classified them as second-degree.

Interestingly, we found that a binomial test indicated that a third-degree relationship could not be rejected for ALM057/ALM079 (p=0.266, Figure 10). However, these two individuals had no common first-or second-degree relatives in the study, and so a third-dgree relationship between could not definitively be definitively shown using additional relatives in a pedigree.

**Figure 10:**
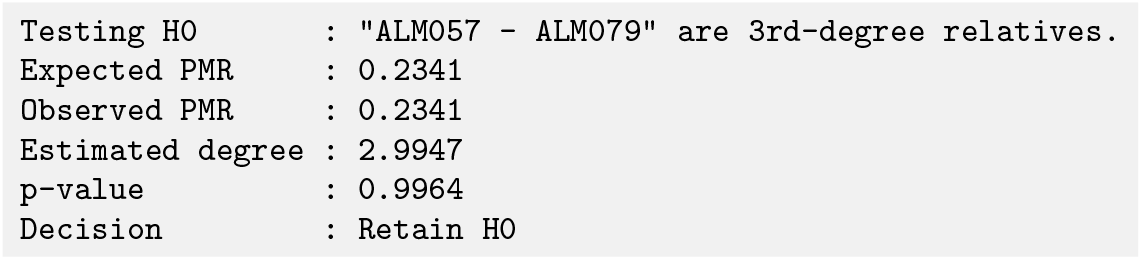
The output from the binomial test for whether the PMR observed for ALM057 and ALM079 is consistent with a third-degree relationship.

For ALM016/ALM017, the observed PMR of 0.2356 was directly in-between what would be expected for second-degree (0.2271) and third-degree (0.2433). However, pedigrees indicated that this was a third-degree relationship, which READ also did not classify correctly. Given that the binomial test rejected both a second- and third-degree relationship, this is likely a pedigree with a higher-level of background relatedness.

Interestingly, ALM016 had an unrelated reproductive partner, ALM015, who was also successfully sequenced. ALM015 and ALM017 were classified as unrelated, but yielded a PMR not inconsistent with a third-degree relationship. Since no individuals were found which filled the pedigree between the generation of the pedigree in which ALM017 lived, and that of ALM015 and ALM016, this inconsistency could explained by one of the missing relatives being somewhat closely-related to ALM016, but not ALM015.

Figure 11 shows the additional diagnostic information available using the BREADR package. ALM057 and ALM079 were identified as unrelated by READ, and second-degree by BREADR. However, the it is also clear from Panel A that the PMR falls somewhere between the distributions for second-degree related and unrelated pairs of individuals.

**Figure 11:**
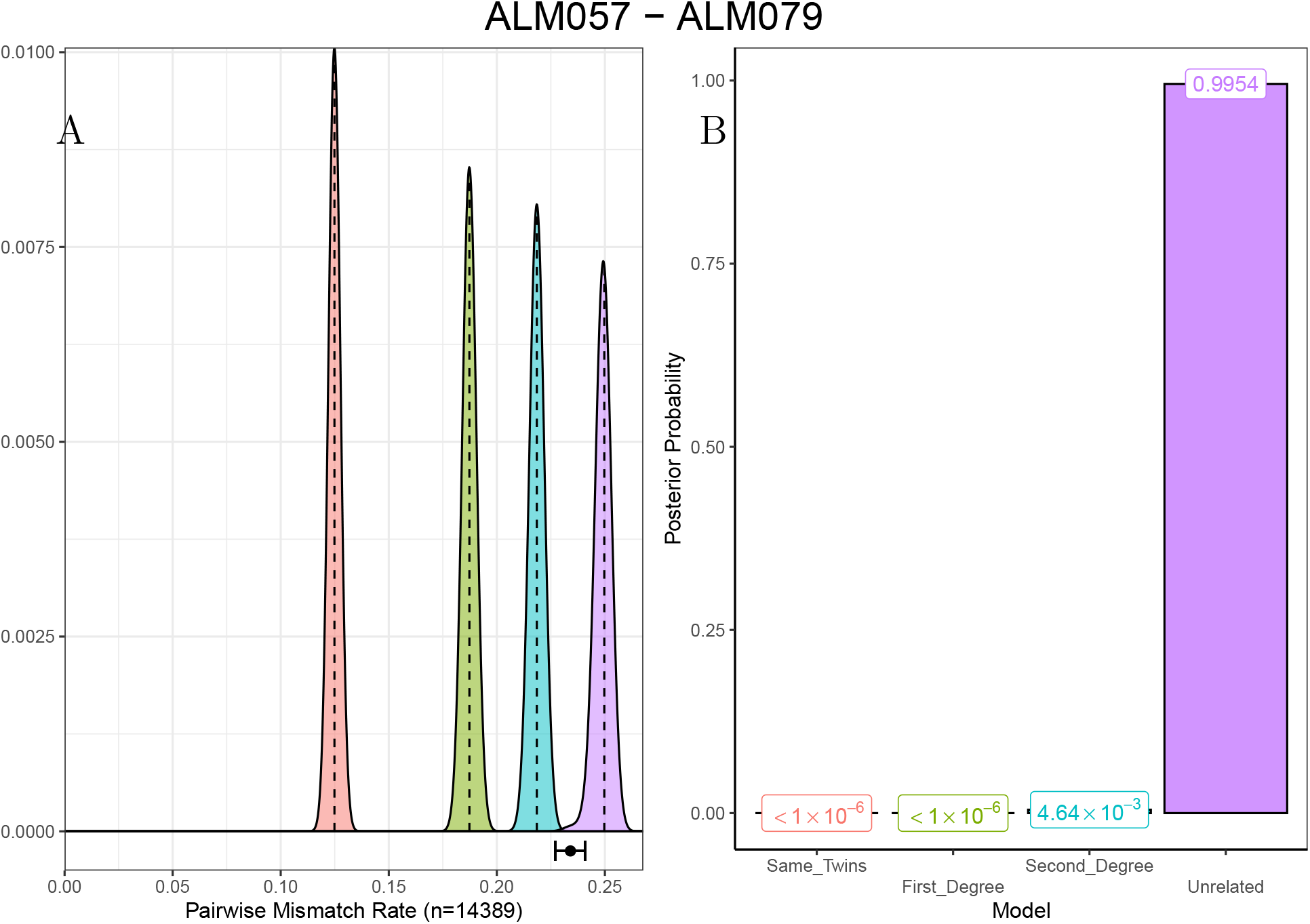
A diagnostic plot for the classification of ALM057/ALM079, with colours indicating classes. Panel A shows the distributions of the pairwise-mismatch rates (PMR) for the same/twins, first-, second-degree, and unrelated classes, with the observed PMR, with 2 standard errors, indicated below. Dashed lines indicated the mode for each class. Panel B shows the normalised posterior probabilities, with values, for all classes.

## 6 Summary and Conclusions

This article describes the usage and functionality of the R package BREADR. This package is used to infer pairwise genetic relatedness between individuals, using the well-understood PMR, up to the second-degree. BREADR can calculate Bayesian posterior probabilities for the classification of genetic relationships on even very low-coverage, pseudohaploid sequence data, in the Eigenstrat data format, common in aDNA studies. Additionally, BREADR provides functionality for plotting the overall genetic relatedness, as well as single plots for specific pairs when a need for exploring the strength of evidence for alternative potential genetic relationships is required. Finally, by allowing for researchers to explore the possibility of other potential levels of genetic relatedness in a statistically rigorous framework, quality control analyses and pedigree reconstructions can be performed with more flexibility.

BREADR has the same limitations of any method that relies on estimating the PMR: namely that estimating the average pairwise-mismatch rate for two unrelated individuals is both difficult to do (and to assess), and is not likely to be constant over large pedigrees. We also identify a need to consider the empirical variability of the expected PMR for *k*-degree relatives that can be observed in large modern datasets. We encourage researchers to consider these limitations when using BREADR, however, as each method has different strengths and weaknesses, we encourage researchers to use multiple methods for estimating genetic relatedness.

We have shown that BREADR performs equally well on both simulated and empirical data, when compared to the peer-reviewed, field-standard PMR-based software READ. However, BREADR is able to assign posterior probabilities to classifications of degrees of relatedness up to the second-degree, or to unrelated. The usefulness for researchers to be able assign statistical confidence to each and every estimated degree of relatedness, even in cases when coverage is extremely low, can not be underestimated when reconstructing a pedigree. We have also shown that when BREADR misclassifies genetic relationships, it is due to expected statistical variation (as READ also misclassified the genetic relationship), or it was due to a true closer genetic relationship (third-degree genetic relationships were misclassified as unrelated). The easy-to-use R-implementation of our method makes BREADR an attractive, and statistically rigorous tool for researchers in archaeogenetics to employ.

## 7 Acknowledgements

We thank Harald Ringbauer and Vincent Braunack-Mayer for important and enlightening discussions regarding the manuscript.

## 8 Bibliography

## Footnotes

1 10.6084/m9.figshare.22574458

2 10.6084/m9.figshare.22574449

3 https://github.com/stschiff/sequenceTools

